# E3 ubiquitin ligase WWP2 regulates stability and tumor suppressive properties of the chromatin remodeler ARID1B

**DOI:** 10.1101/2025.03.01.640953

**Authors:** Pradipta Hore, Sandipkumar Bambhaniya, Murali Dharan Bashyam

**Affiliations:** Laboratory of Molecular Oncology, BRIC-Centre for DNA Fingerprinting and Diagnostics, Hyderabad, Telangana, India; Graduate studies, BRIC-Regional Centre for Biotechnology, Faridabad, India

**Keywords:** ARID1B, WWP2, Ubiquitination, Proteasomal degradation, Tumor suppression

## Abstract

ARID1B, a key subunit of the SWI/SNF (also known as BAF) chromatin remodeling complex, is characterized as a canonical tumor suppressor across various cancer types. Although the downregulation of ARID1B transcript levels has been observed in many cancers, its regulation at the protein level is comparatively less studied. Here, we identify WWP2, an E3 ubiquitin ligase, as a novel interacting partner of ARID1B. Our results show that using its WW domains, WWP2 interacts with the PPxY motif located at the N-terminus of ARID1B. We further demonstrate that wild-type WWP2, but not its catalytically inactive mutant, regulates ARID1B protein stability through ubiquitination-mediated proteasomal degradation. Interestingly, WWP2 appears to facilitate non-canonical K27- and K29-linked polyubiquitination of ARID1B. Additionally, silencing WWP2 expression results in a decrease in ubiquitination and a subsequent increase in ARID1B protein levels, indicating that WWP2 plays a crucial role in regulating ARID1B stability. Finally, based on several tumorigenic assays performed in cell lines and mouse xenograft models, we show that WWP2 may modulate ARID1B-mediated tumor suppression. Our results therefore highlight a novel mechanism of post-translational regulation of ARID1B, which may have implications in ARID1B-mediated tumor suppression.

## Introduction

The mammalian SWItch/Sucrose Non-Fermentable (SWI/SNF) or BRG1/Brm associated factor (BAF) is a multi-subunit ATP-dependent chromatin remodeling complex involved in several cellular processes. The complex is categorised into three forms: canonical (cBAF), Polybromo (PBAF), and non-canonical (ncBAF) or GLTSCR1-like containing (GBAF)[1]. The AT-Rich Interaction Domain 1B (ARID1B) also called BAF250b, along with its paralog ARID1A, are mutually exclusive components of the cBAF complex [2]. ARID1B is a 250kDa protein containing multiple structural units including a 94-amino acid-long ARID domain, multiple LXXLL motifs that are expected to facilitate protein-protein interactions, a C-terminal Armadillo (ARM) repeat-containing BAF250_C domain that aids in BAF complex formation, and a nuclear localisation signal (NLS) (Supp. Fig. 1A)[3–5]. Majority of the protein is constituted by a classical intrinsically disordered region (IDR) (Supp. Fig. 1A) [6]. ARID1B has been reported to play important roles in fundamental cellular processes such as transcriptional regulation [7], mRNA splicing [8], DNA repair [9], and cell proliferation [10]. Multiple studies have revealed the regulation of *ARID1B* through mutations [11], chromosomal rearrangements [12], and epigenetic modifications such as promoter DNA methylation [13] with implications for tumorigenesis. However, the mechanism of ARID1B regulation at the protein level is largely unknown.

Ubiquitination is a common reversible post-translational protein modification majorly classified as mono or poly-ubiquitination [14]. Ubiquitin, a 76 amino acids long protein, uses either of seven lysine residues (K6, K11, K27, K29, K33, K48, and K63) or the α-amino group of the N-terminal methionine to effect linear or branched chain poly-ubiquitination with different combinations [15]. The ubiquitination process occurs in three steps: first, an ‘E1’ enzyme activates the ubiquitin molecule which is subsequently transferred to an ‘E2’ enzyme. The E2 interacts with the ubiquitin ligase ‘E3’ which in turn utilizes the C-terminal glycine of ubiquitin to link to the ε-amino group of a lysine residue on the substrate protein [16,17].

E3 ubiquitin ligases are categorized into three types according to their E2 binding domain structure and ubiquitin transfer mechanism: RING (Really Interesting Gene), HECT (homologous to the E6-associated protein carboxyl domain), and RBR (RING-between-RING) [18]. WW domain-containing E3 ubiquitin protein ligase 2 (WWP2), which belongs to the NEDD4 family of HECT ubiquitin ligases [19], is well characterized with respect to its role in transcription [20], cell proliferation [19], and DNA repair [21]. WWP2 is a ∼100 kDa protein that mainly comprises three domains: an N-terminal C2 domain, four WW domains responsible for the recognition of Proline-rich motifs on the substrates, and a C-terminal catalytic HECT domain responsible for the ubiquitin ligase activity [22]. *WWP2* is classified as an oncogene with well-established roles in hepatocellular [23,24], oral [25], breast [26], and prostate [26] carcinomas, as well as in glioma [27]. Previous studies have highlighted the involvement of WWP2 in regulating multiple cancer genes such as *PTEN* [19], *OCT4* [20], etc.

Here, we report a novel interaction between ARID1B and WWP2. We further show that ARID1B stability **is** regulated by WWP2 and that WWP2 appears to modulate ARID1B-mediated tumor suppression.

## Results

### ARID1B interactome reveals possible involvement in multiple cellular processes

To understand the possible mechanism of ARID1B regulation at the post-translational level, we examined its interactome using affinity purification of Halo-tagged ARID1B followed by LC-MS/MS (Liquid Chromatography-Tandem Mass Spectrometry) in HEK293T cells (Fig. 1A). We also mined an ARID1B interactome published earlier [8]. A combined robust analysis of all screens revealed a common list of 180 putative ARID1B interacting proteins (Supp. Fig. 1B, Supp. Table 1). Gene ontology (GO) analysis revealed the ARID1B interactome to be enriched in fundamental cellular processes including G0 to G1 transition, nucleotide excision repair, and cytoplasmic translation (Fig. 1B), in addition to previously identified roles in nucleosome binding, transcriptional regulation, and RNA splicing (Fig. 1C). Interestingly, scrutiny of the list of interacting proteins revealed only two E3 ubiquitin ligases, both belonging to the HECT family, namely WWP2 and its paralog WWP1 (Fig. 1D).

**Fig. 1:**
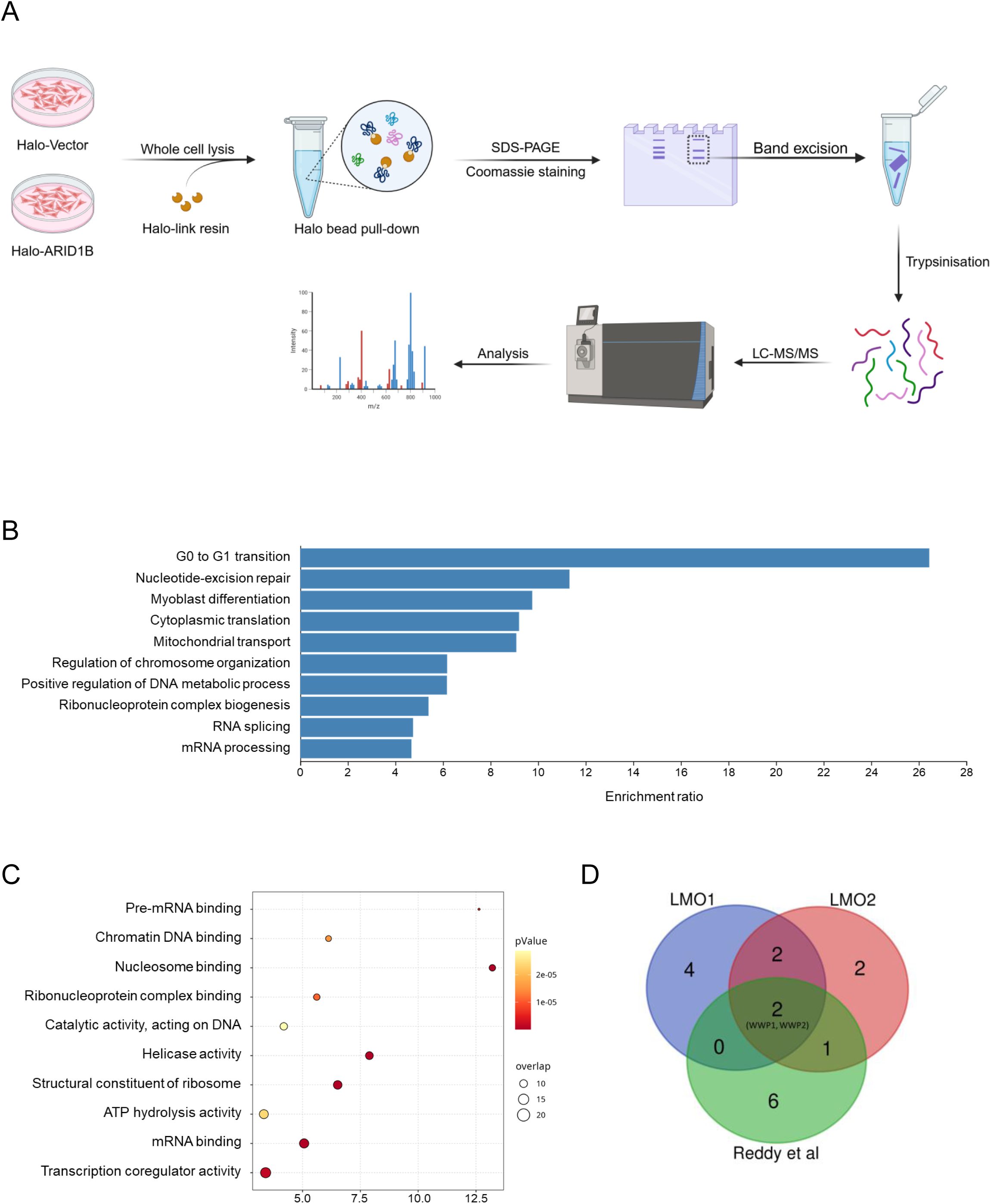
The ARID1B interactome reveals its involvement in multiple cellular processes. Schematic representation of methodology to determine the ARID1B interactome (A). Gene Ontology (GO) analysis of the ARID1B interactome for ‘cellular processes’ (B) and ‘molecular function’ (C). Ubiquitin ligases interacting with ARID1B identified from multiple interactome lists (D).

### ARID1B interacts with E3 ubiquitin ligases WWP1 and WWP2

To validate our affinity purification results, we tested the interaction of ARID1B with WWP1 and WWP2. Ectopically expressed ARID1B interacted specifically with endogenous as well as ectopically expressed WWP1 and WWP2 in HEK293T (Fig. 2A and 2B). We further confirmed the existence of the ARID1B-WWP2 complex by demonstrating that WWP2 co-immunoprecipitated with endogenous ARID1B in HCT116 cells (Fig. 2C). We generated multiple deletion constructs of SFB-tagged WWP2 (Fig. 2D) to map its ARID1B-binding region. Affinity precipitation results suggested that WW domains on WWP2 could be important for the interaction, as reported earlier [28] (Fig. 2E). Similarly, to map the binding site of WWP2 on ARID1B, we used a series of SFB-tagged ARID1B carboxy-terminal deletions generated previously in the lab (Fig. 2F). We co-expressed each of these deletion constructs separately with full-length WWP2 or WWP1. The affinity purification results confirmed that both WWP2 (Fig. 2G) and WWP1 (Supp. Fig. 2) may interact with the N-terminal region of ARID1B. We further verified this result by assessing the interaction with endogenous WWP2 (Fig. 2H).

**Fig. 2:**
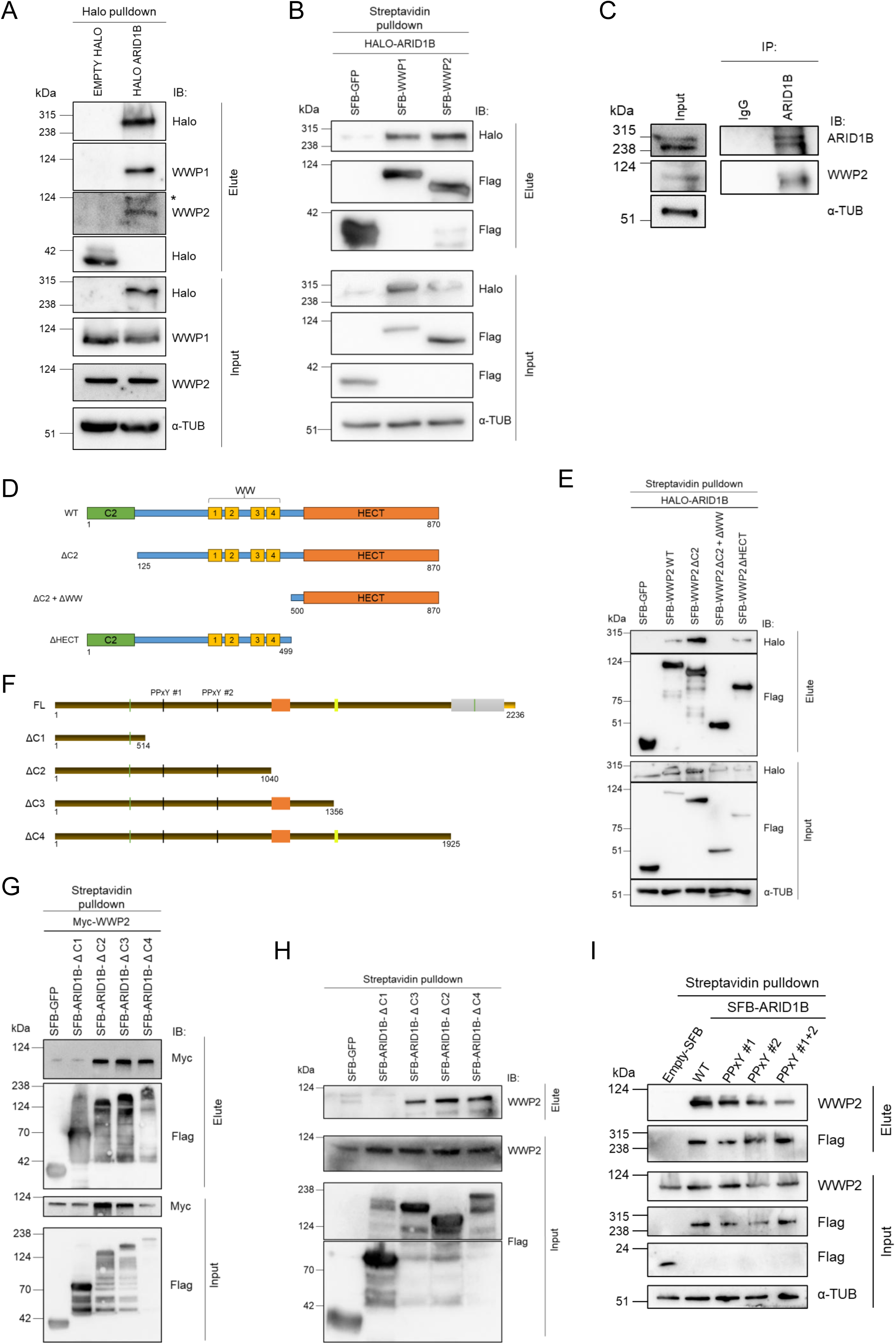
WWP1 and WWP2 are novel interacting partners of ARID1B. Affinity-based pull-down to validate the interaction of Halo-tagged ARID1B with endogenous (A) and ectopically expressed (B) WWP1 and WWP2 in HEK293T cells. * indicates non-specific band. Immunoprecipitation of endogenous ARID1B, validating the interaction with WWP2 in HCT116 cells (C). Schematic representation of the full-length and various deletion mutants of WWP2 (D) and assessment of their interaction with ARID1B (E). Schematic representation of the full-length and various C-terminal deletion mutants highlighting the location of PPxY motifs in ARID1B (F) and assessment of their interactions with ectopically expressed (G) and endogenous (H) WWP2. Interaction between endogenous WWP2 and wild-type or various PPxY mutated derivatives of ARID1B (I). All experiments were performed at least in three biological replicates; representative images are shown.

The WW domain, typically 35-40 amino acids long, interacts with proline-rich motifs such as PPxY, PPLP, and phosphorylated serine/threonine-proline sites via its conserved tryptophan residues [29]. We detected **two** such PPxY motifs (PPxY#1 located at 523-526 aa, and PPxY#2 located at 785-788 aa) (Fig. 2F) through ARID1B sequence analysis. We used site-directed mutagenesis to mutate these two sites in ARID1B and further evaluated interaction with WWP2. Our results revealed a significant reduction of interaction between PPxY mutant ARID1B and WWP2 (Fig. 2I).

### WWP2 regulates ARID1B protein level by facilitating proteasomal degradation

To characterize the functional significance of the interaction between ARID1B and WWP1/WWP2, we evaluated ARID1B levels in cells upon ectopic expression of WWP2. The presence of increasing amounts of wild-type (but not the catalytically inactive mutant (WWP2^C838A^)) WWP2 led to a reduction in ectopically expressed (Fig. 3A and 3B) and endogenous (Fig. 3C) ARID1B levels without affecting its localisation (Supp. Fig. 3A), negating a possibility of non-canonical effect of ubiquitination. Similar results were obtained with wild-type and catalytically inactive mutant WWP1^C890S^ (Supp. Fig. 3B). On the other hand, we observed an elevation in endogenous ARID1B protein levels upon shRNA-mediated stable knockdown of WWP2 (Fig. 3D). These results were further substantiated by detecting elevated transcript levels of ARID1B target genes *CDKN1B* and *TP53* upon WWP2 knockdown (Supp. Fig. 3C). Further, cycloheximide chase experiments revealed that the downregulation of WWP2 caused a significant increase in the stability of ectopically expressed ARID1B (Fig. 3E and 3F). Similarly, we observed a significant reduction in ARID1B half-life in the presence of wild-type but not the catalytic mutant form of WWP2 (Fig. 3G and 3H) or WWP1 (Supp. Fig. 3D and 3E). Given that WWP2 is a known HECT-domain-containing E3 ligase involved in the ubiquitin-mediated proteasomal degradation of its substrates, it is expected to regulate ARID1B levels in a similar manner. Indeed, treatment with the proteasome inhibitor MG132 caused a significant elevation in ARID1B protein levels in HCT116 cells expressing wild-type WWP2, whereas no such increase was observed in cells expressing the catalytic mutant WWP2 or the vector control (Fig. 3I). Next, we ectopically expressed wild-type and catalytically inactive WWP2 under conditions of WWP2 stable knockdown. ARID1B endogenous protein level was partially rescued by wild-type (but not mutant) WWP2 (Fig 3J).

**Fig. 3:**
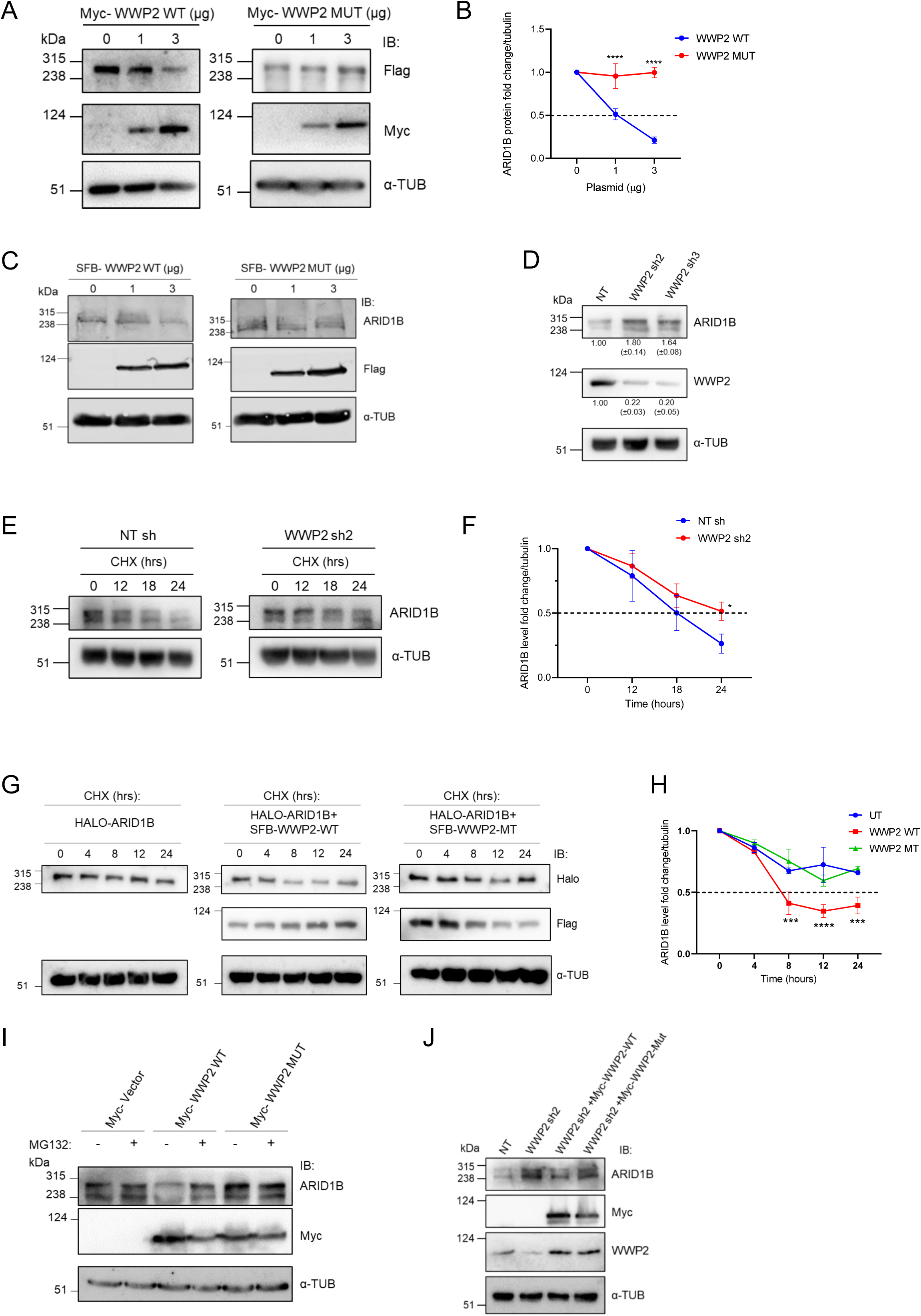
ARID1B is subjected to WWP2-mediated regulation via proteasomal degradation. Assessment of ectopically expressed ARID1B levels when exposed to increasing amounts of wild-type (left) or catalytically inactive mutant (right) of WWP2 in HEK293T cells (A); quantification of change in ARID1B protein levels is shown in panel B. Change in endogenous ARID1B protein levels under similar conditions as (A) is shown in panel (C). Evaluation of ARID1B protein levels upon shRNA-mediated knockdown of WWP2 in HEK293T (D). Protein intensity fold change is represented as mean ± SD. Assessment of endogenous ARID1B levels by cycloheximide chase assay upon shRNA-mediated down-regulation of WWP2 (E); quantitation of change in ARID1B levels is shown in panel F. Evaluation of ARID1B protein stability by cycloheximide chase assay in the presence or absence of wild-type or mutant WWP2 (G); quantitation of change in ARID1B levels is shown in panel H. Evaluation of change in ARID1B protein levels in the presence of wild-type or mutant WWP2 upon MG132 treatment in HEK293T cells (I). Change in ARID1B endogenous protein level was evaluated upon ectopic expression of wild-type and catalytically inactive mutant of WWP2 in WWP2 knockdown HCT116 cells (J). All experiments were performed at least in three biological replicates; representative images are shown. All quantifications (panels B, F and H) are from three independent experiments. EV, empty vector; WT, wild-type; MT, mutant; UT, un-transfected. All statistical analyses were performed using two-way ANOVA (N=3), \**P*<0.05; \*\**P*<0.01; \*\*\**P*<0.001; \*\*\*\**P*<0.0001.

### WWP2 facilitates K27/K29 linked polyubiquitin chain elongation on ARID1B

Further to investigate whether WWP2 mediated ARID1B proteasomal degradation occurs via ubiquitination of the latter, we performed affinity precipitation of ARID1B under denaturing condition in the presence of MG132 to detect ubiquitination level. We observed that ARID1B ubiquitination levels were elevated by wild-type (but not mutant) WWP2 (Fig. 4A). This result was further supported by a reduction in the levels of ARID1B ubiquitination observed upon WWP2 knockdown in HEK293T (Fig. 4B) as well as HCT116 (Fig. 4C) cells. Interestingly, a ubiquitination assay using various ubiquitin mutants revealed that WWP2 appeared to facilitate K27 (Lysine-27) and K29 (Lysine-29)-linked polyubiquitin chain formation on ARID1B (Fig. 4D), which is in contrast to the previously reported model of preferential K63 (Lysine-63) poly-Ubiquitin chain formation by HECT family ligases [30,31]. To further validate the result, we performed ARID1B ubiquitination assay in the presence of ubiquitin mutants namely K27R, K29R, K48R, or K0 separately. In agreement with the previous results, ARID1B ubiquitination was markedly reduced in cells expressing the K27R or K29R ubiquitin mutants, whereas significant ubiquitination was observed with the K48R mutant (Fig. 4E), indicating a novel K27/K29 linked polyubiquitin chain elongation by WWP2. Finally, based on ubiquitin lysine-linkage specific antibodies, we confirmed specific K27 (but not K63)-linked chain elongation on ARID1B in ubiquitination assays (Fig. 4F, Supp. Fig. 4A).

**Fig. 4:**
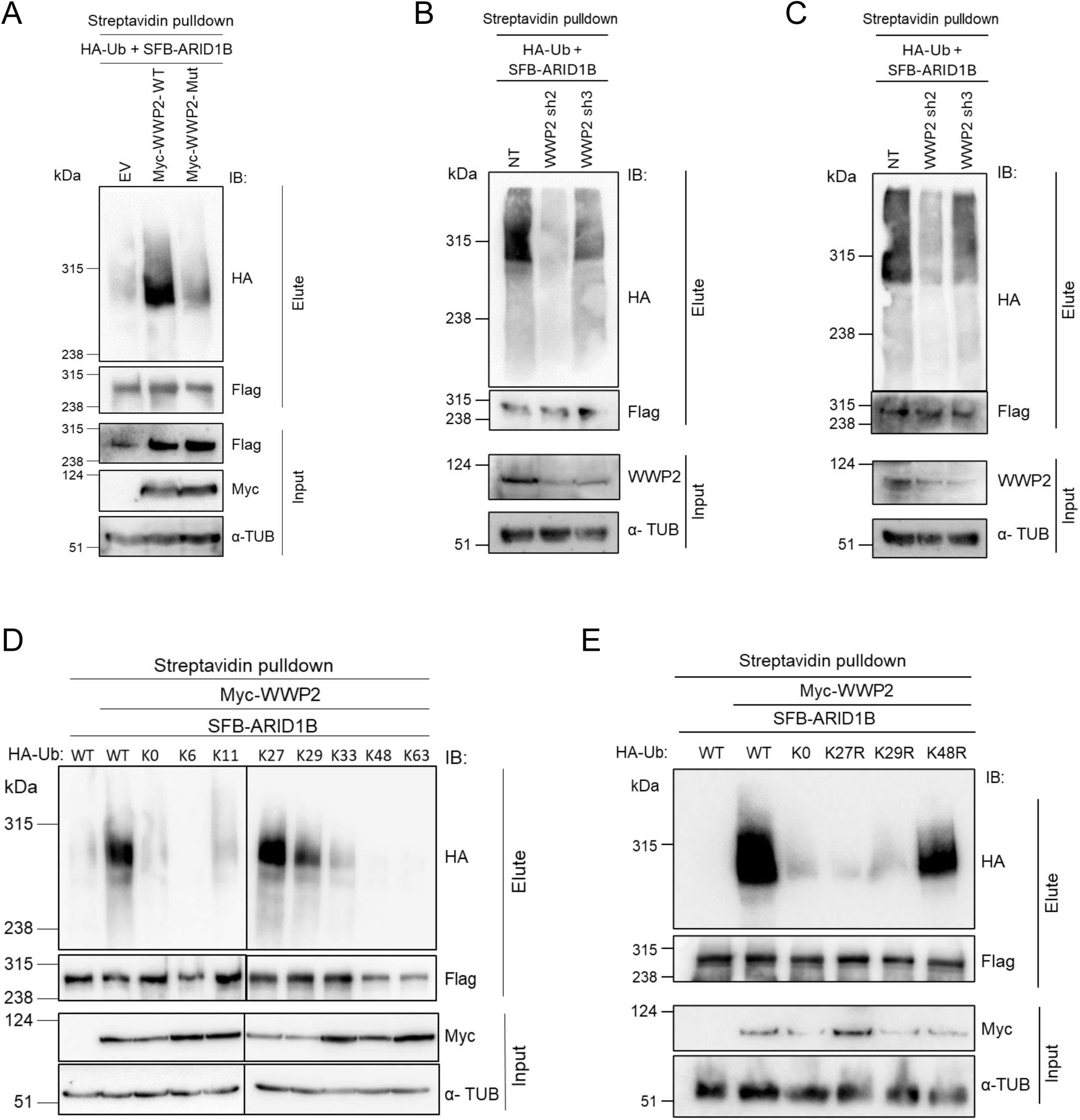
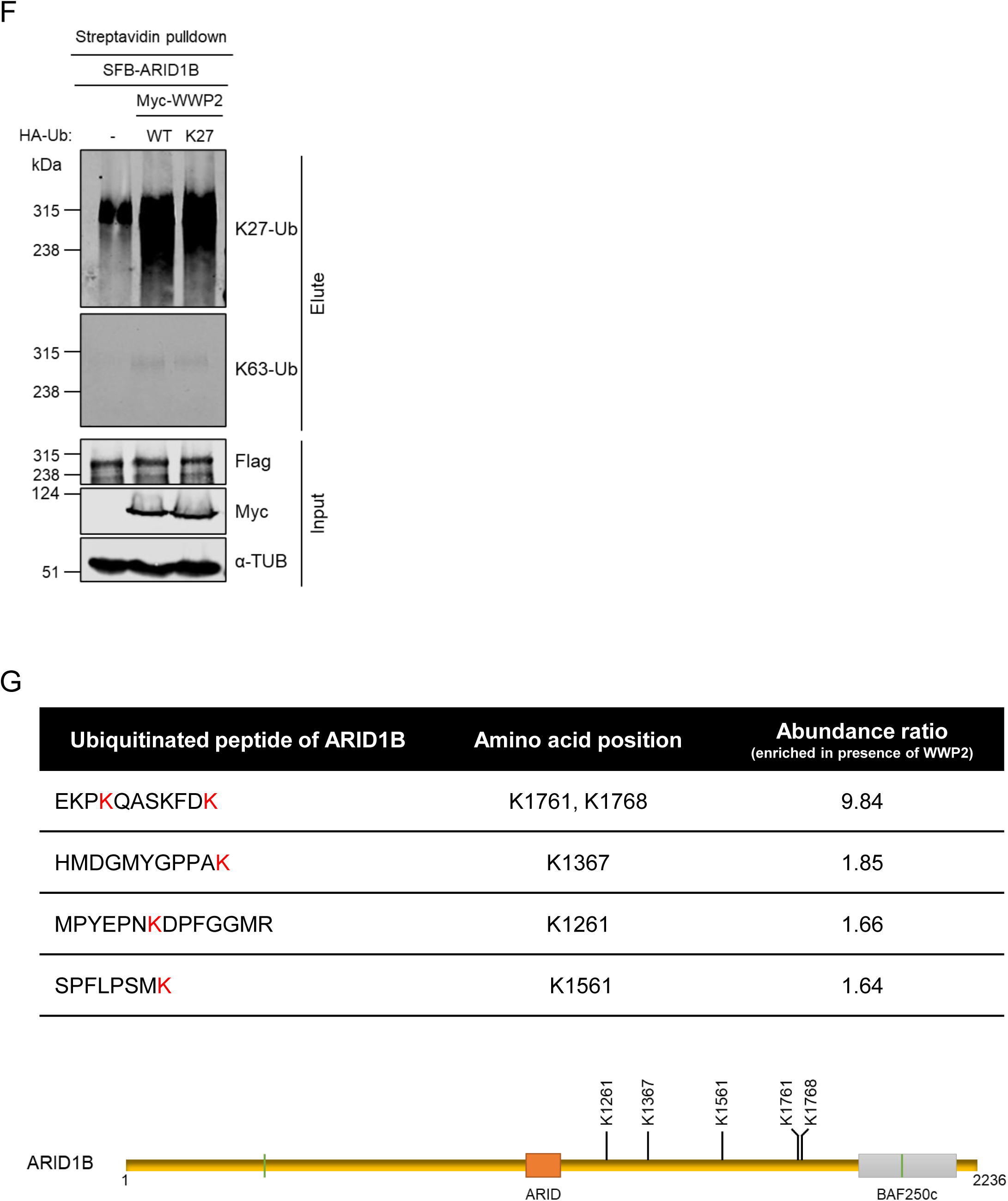
WWP2 performs K27/K29 linkage specific polyubiquitination on ARID1B. Evaluation of ARID1B ubiquitination in the presence of ectopically expressed wild-type or mutant WWP2 in HEK293T cells (A) or upon shRNA-mediated knockdown of WWP2 in HEK293T (B) and HCT116 (C) cells. Assessment of specific poly-ubiquitin chain formation of ARID1B in the presence of wild-type WWP2 and wild-type or various lysine mutant constructs of Ubiquitin (D, E). Analysis of linkage-specific polyubiquitin chain elongation of ARID1B in the presence of WWP2, upon immunoblotting with endogenous K27-linkage and K63-linkage specific ubiquitin antibodies (F). List of ubiquitinated lysine residues of ARID1B (top) and respective positions on ARID1B (bottom) detected to be enriched by mass spectrometry analysis in HEK293T cells (G). All experiments were performed at least in three biological replicates; representative images are shown.

To identify the specific ARID1B lysine sites modified by ubiquitination, ARID1B and ubiquitin (Ub) were co-expressed in HEK293T cells either in the presence or absence of ectopically expressed wild-type WWP2, followed by Streptavidin-agarose bead-based affinity precipitation and mass spectrometry (MS) analysis (Supp. Fig. 4B). Five ubiquitinated lysine residues of ARID1B were detected to be enriched in the presence of WWP2 (Fig. 4G, Supp. Fig. 4C).

### WWP2 potentiates ARID1B depletion-dependent tumorigenic features

Having established a critical role of WWP2 in regulating ARID1B protein levels, and given the previously established oncogenic and tumor suppressor functions of WWP2 and ARID1B, respectively, we attempted to explore the role of WWP2 in ARID1B downregulation-dependent tumor progression. To this end, we first generated ARID1B knockdown in HCT116 cells and validated its tumor suppressor function using various assays; there was a significant elevation in several tumorigenic features following a reduction in ARID1B levels (Supp. Fig. 5A-D), as previously reported [10,13]. We subsequently generated WWP2 knockdown in HCT116 cells already possessing knockdown of ARID1B, which caused a reduction in the elevated tumorigenic phenotypes resulting from ARID1B knockdown (Fig. 5A-D, Supp. Fig. 5E-G). Finally, we evaluated the tumorigenic potential of HCT116 cells having either single ARID1B knockdown or combined ARID1B+WWP2 knockdown in Foxn1^−/−^ nude female mice. Consistent with our findings from cell-line based tumorigenesis assays, concurrent silencing of ARID1B and WWP2 resulted in a pronounced reduction in tumor growth rate compared to single ARID1B knockdown (Fig. 6A-C), supporting a possible modulation of ARID1B tumor suppressor function by WWP2.

**Fig. 5:**
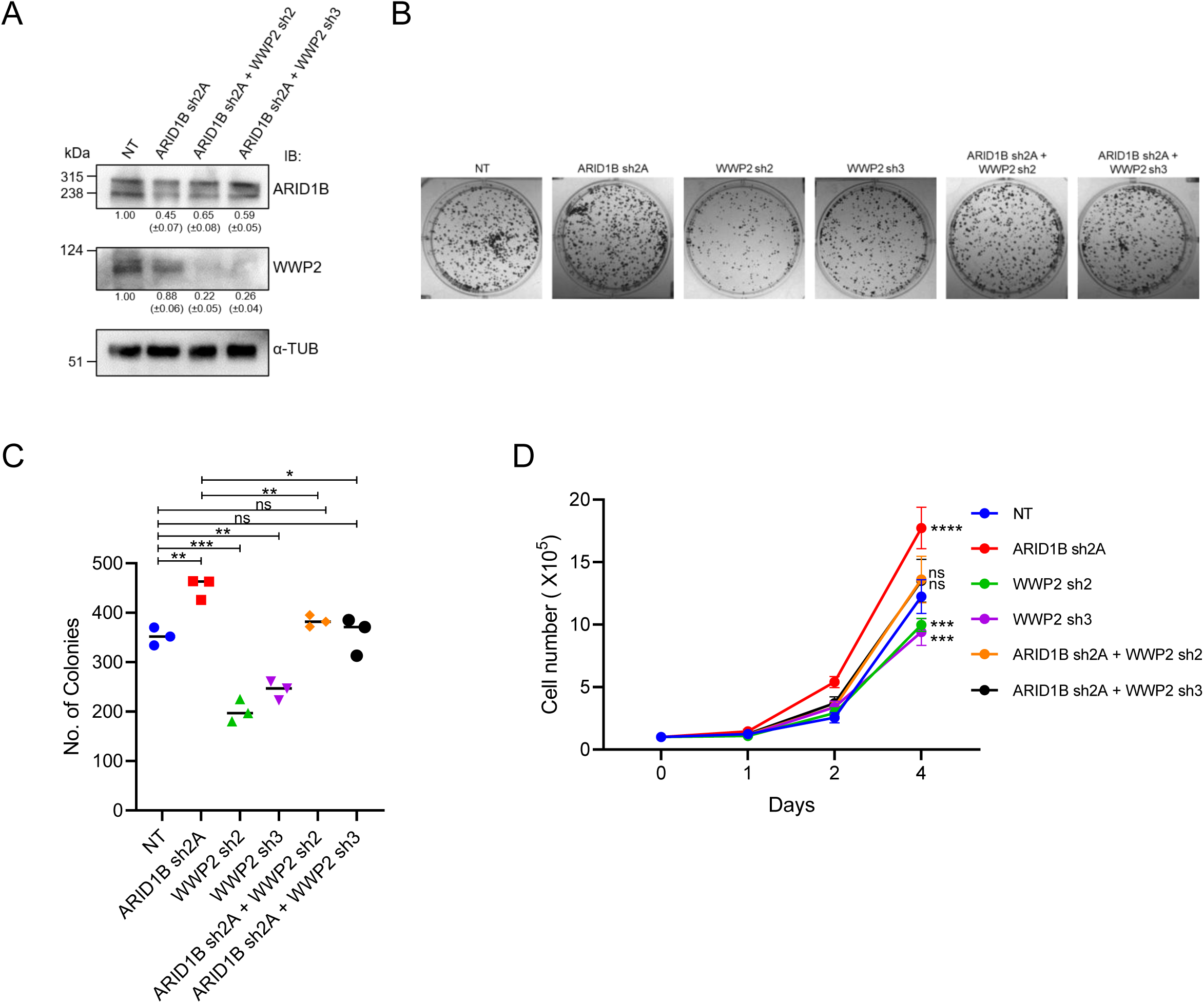
WWP2 modulates ARID1B tumor suppressor properties. shRNA mediated knockdown of ARID1B (sh2A) and subsequently of WWP2 (sh2 or sh3) in HCT116 cells (A). Colony formation (B, C), and cell growth (D) assays were performed with various combinations of down regulation of ARID1B and WWP2, as indicated. All results are from three experiments and presented as mean ± SD. All experiments were performed at least in three biological replicates; representative images are shown. All statistical analyses were performed using either unpaired Student’s T test or two-way ANOVA. \**P*<0.05; \*\**P*<0.01; \*\*\**P*<0.001; \*\*\*\**P*<0.0001; ns, not significant

**Fig. 6:**
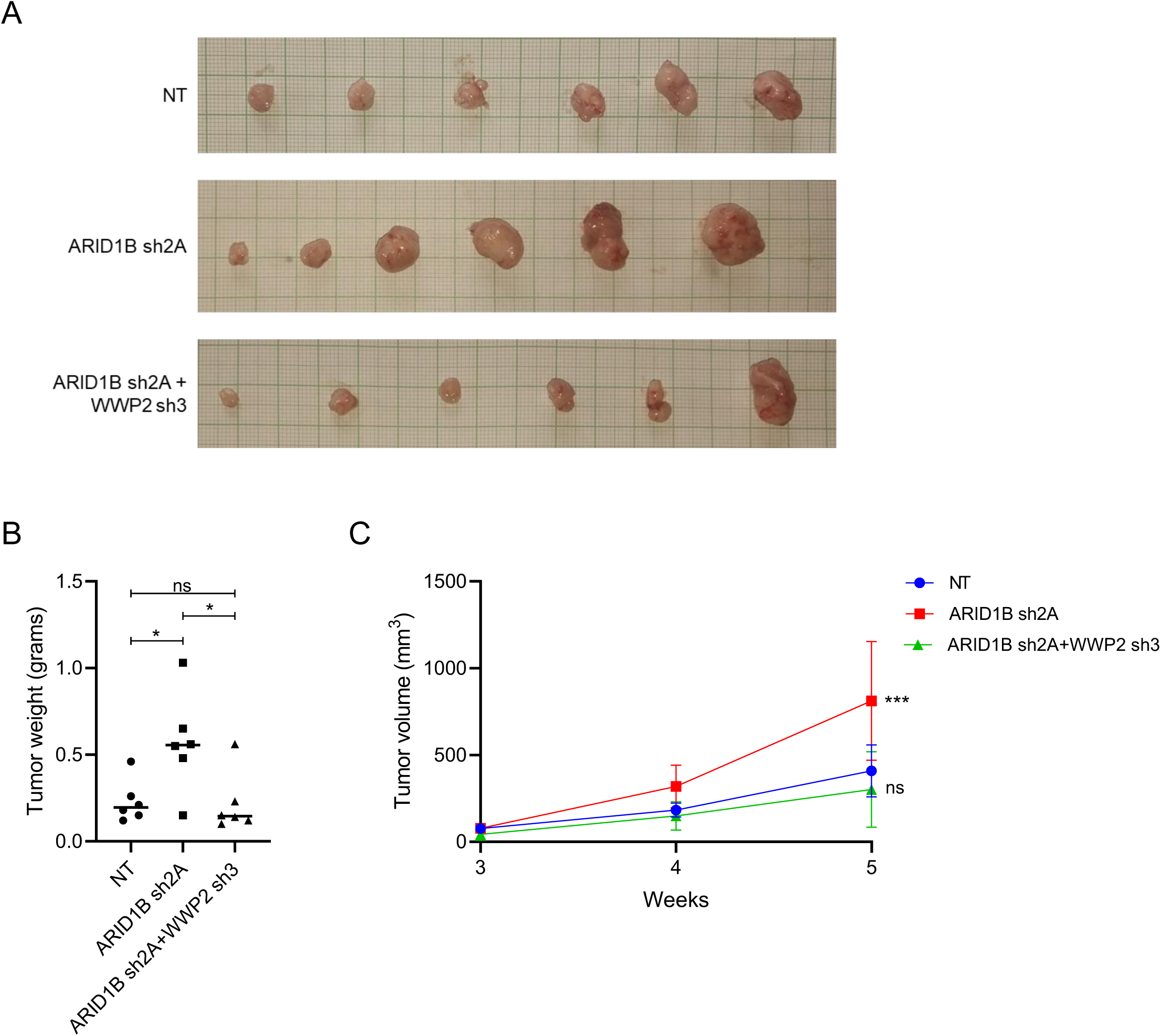
WWP2 promotes tumor growth in a mouse xenograft model by downregulation of ARID1B. The tumor formation ability of single ARID1B or dual ARID1B+WWP2 knockdown HCT116 cells were evaluated in Foxn1^-/-^ nude mice. Representative tumor images are shown in panel A. Quantification of tumor weight and of volume over time (mean for 6 animals) are shown in panels B and C, respectively. All statistical analyses were performed using either unpaired Student’s T test or two-way ANOVA. \**P*<0.05; \*\**P*<0.01; \*\*\**P*<0.001; \*\*\*\**P*<0.0001; ns, not significant

**Fig. 7:**
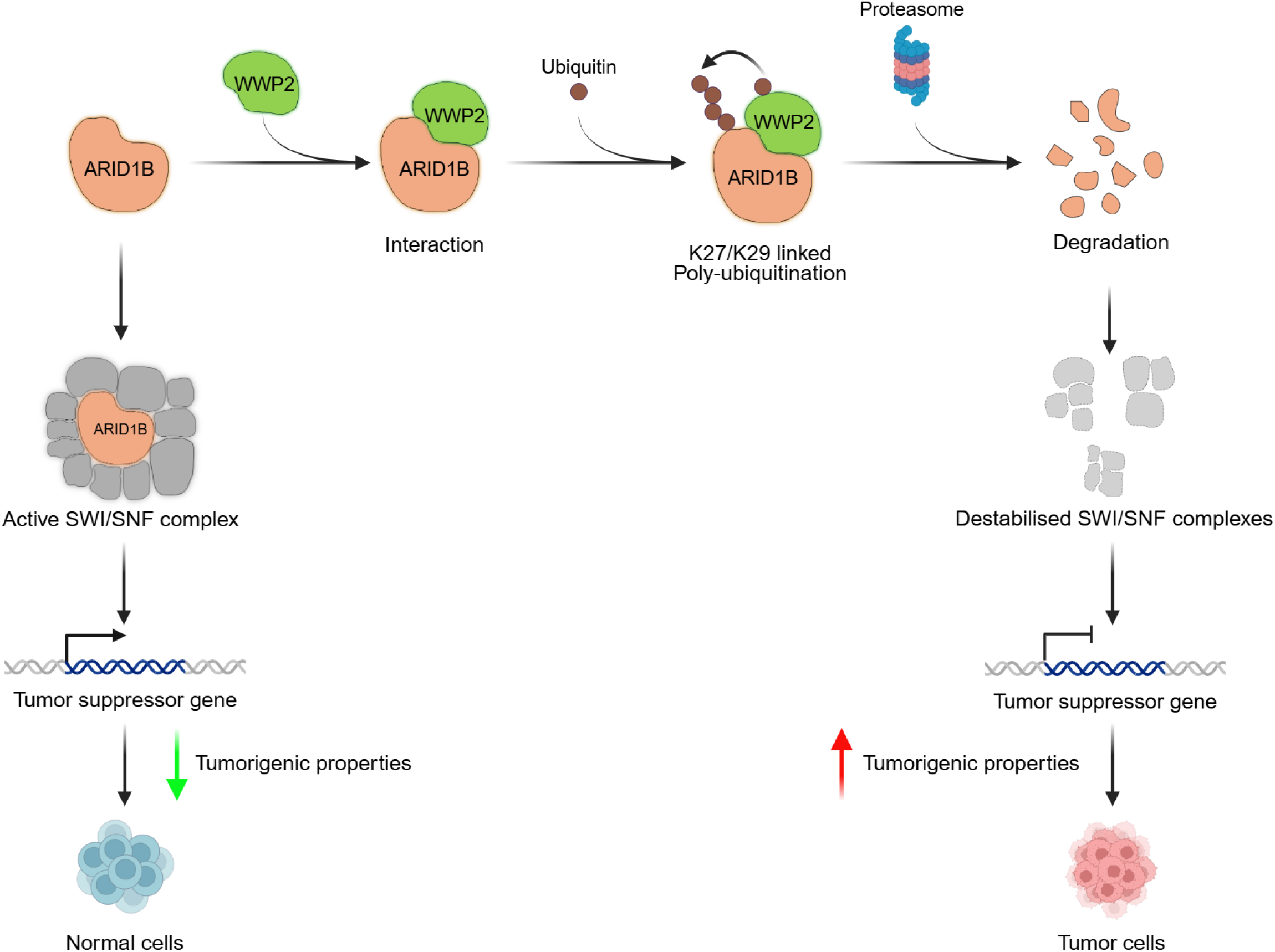

## Discussion

In the current study, we attempted to determine modes of regulation of ARID1B, an important chromatin remodeler, at the protein level. The scrutiny of ARID1B-interactome, generated in-house and elsewhere, revealed HECT family ubiquitin ligases WWP1 and WWP2 as novel interactors of ARID1B. Our results revealed ARID1B N-terminal PPxY motifs spanning amino acids 437-1035 to be involved in interaction with WWP1/WWP2 suggesting a complete non-overlap with the ARID1B region (C-terminal BAF250_C domain) responsible to form the BAF complex [32]. Interestingly, mutating both PPxY motifs within this region reduced but did not entirely disrupt ARID1B’s interaction with WWP1/P2, suggesting that additional proline-rich motifs in ARID1B could contribute to binding with the WW domains of WWP1/WWP2.

Previous reports showed that E3 ubiquitin ligases WWP1 and WWP2 shared multiple common substrates like PTEN [19,33], Dvl2 [34,35], and Atophin-1 [36]. Our results indicate ARID1B to be yet another common substrate for these two E3 ligases. WWP1 and WWP2 have been shown to form heterodimers and regulate cellular homeostasis of p73 and ΔNp73 [37]. While we confirmed the independent action of WWP1 and WWP2 on ARID1B, further studies should focus on deciphering the detailed mechanism of WWP1 dependent ARID1B regulation and determining its functional consequences for transcriptional control and tumorigenic properties. Interestingly, WWP2 is known to regulate another SWI/SNF complex component, Srg3 (mouse homolog of human BAF155/SMARCC1) [38]. Further, WWP2 is shown to utilize different ubiquitination chains including K11, K27, K48, and K63, to perform poly-ubiquitination [39]. Of note, K63-linked ubiquitination of Dvl2 by WWP2 was shown to be required for the former’s ability to form condensates [35]. It will be interesting to determine whether WWP2 independent ubiquitination of ARID1B has any effect on its phase separation properties [6]. Previously, ARID1B has been reported to be **a** part of **a** ubiquitin ligase system containing Cul2, EloC and Roc1 responsible for histone H2B monoubiquitination. Mutation in the BC box motif in ARID1B results in auto-ubiquitination followed by proteasomal degradation by the same complex [40]. Our study highlights an independent mode of post-translational regulation of ARID1B that probably perturbs its tumor suppressor function.

Among the various Lysine residues that participate in polyubiquitination, K48- and K63-linked polyubiquitination are the most extensively studied. K48-linked polyubiquitination is primarily associated with proteasomal degradation, whereas K63-linked polyubiquitination is non-degradative and predominantly involved in mediating protein interactions and activating signaling pathways [15,41]. Recent research has highlighted the role of K11- and K27-linked polyubiquitination in ubiquitin-mediated proteasomal degradation [42–44]. Although HECT E3 ubiquitin ligases are classically thought to catalyse K48- or K63-linked ubiquitin chain elongation [30,31,45], several members of this family, including WWP1, ITCH, and HACE1, have recently been shown to mediate K27- and K29-linked polyubiquitination as predominant chain types [46–48]. In this study, we demonstrated that WWP2 mediates non-canonical K27 and K29-linked polyubiquitination of ARID1B, resulting in its proteasomal degradation. In other studies, WWP2 has been reported to form a complex with another HECT E3 ubiquitin ligase, ITCH, to mediate the K27-linked poly-ubiquitination of SHP-1 phosphatase [49]. Interestingly, ITCH was identified in only one of our ARID1B interactomes (LMO1) but could not be independently validated using pull-down followed by immunoblotting (Supp. Fig. 6). Recent studies showed that K27-linked poly-ubiquitination is predominantly a nuclear modification and plays an important role in cell proliferation, DNA repair, and cell cycle progression, besides proteasomal degradation [50]. Further studies are required to determine whether ARID1B K27-ubiquitination is related to these processes. K29-linked polyubiquitination has also been associated with the regulation of protein levels [51].

ARID1B has been extensively studied for its critical role in tumor suppression, presumably due to its function in transcriptional activation of classical tumor suppressors, including *CDKN1A* (p21), *CDKN1B* (p27), and *TP53* (p53) [5,52] and ARID1B downregulation has been linked to oncogenic stimulation [10,13]. Here, we demonstrated that the oncogenic properties associated with ARID1B downregulation can be potentially reversed by concomitant downregulation of WWP2. This discovery is important in the context of recent reports highlighting the utility of targeting synthetic lethal genes linked to BAF complex components in several cancers [53,54]. WWP2 could thus potentially represent a promising therapeutic target for addressing ARID1B-mediated tumorigenesis. To summarise, this study reveals a novel mechanism of non-canonical polyubiquitination-mediated post-translational regulation of ARID1B by the ubiquitin ligase WWP2. WWP2 upregulation decreases ARID1B protein levels, promoting tumorigenic phenotypes.

## Methods

### Cell Culture and Modifications

HEK293T (a kind gift from Dr. Rashna Bhandari, BRIC-Centre for DNA Fingerprinting and Diagnostics (BRIC-CDFD), Hyderabad, India) and HCT116 cells (obtained from BRIC-National Centre for Cell Science (BRIC-NCCS), Pune, India) were cultured as described previously [5].

For generating knockdowns, ARID1B short hairpin RNAs (shRNA; Clone IDs TRCN0000420576 (sh2A) and TRCN0000107360 (sh3A)) and WWP2 shRNAs (5′-ATAAAGCGAAAGTAGGTGAGG-3′ (sh2); 5′-TTGACGATTATGCACCTTGGG-3′ (sh3)) were co-transfected with lentiviral packaging plasmids VSV.G and psPAX2 into HEK293T cells using polyethyleneimine (Sigma-Aldrich, St. Louis, MO, USA). Cell lysis was performed following **a** previous protocol [5]. The extent of knockdown was determined by immunoblot analysis with specific antibodies. Tumorigenic assays including cell growth, viability (3-[4,5-dimethylthiazol-2-yl]-2,5 diphenyl tetrazolium bromide (MTT)), colony formation, and transwell migration assays were performed as described previously [5].

### Antibodies, Immunoblotting, and Immunocytochemistry

Antibodies - WWP1 (Cat # ab43791), ARID1B (Cat # ab57461), and K27-linkage specific Ubiquitin (Cat # ab181537) (Abcam, Cambridge, UK). Hemagglutinin (Cat # H6908), GAPDH (Cat # G8795), Myc clone 9E10 (Cat # M4439), Flag (Cat # F1804), and α-Tubulin (Cat # T6074) (Sigma-Aldrich, MO, USA). Halo (Cat # G9211) (Promega Corporations, Madison, WI, USA). ITCH (Cat # A8624), ARID1B (Cat # A15488) (Abclonal, MA, USA). WWP2 (Cat # A302-936A), K63-linkage specific Ubiquitin (Cat # 14-6077-82), and all HRP-conjugated secondary antibodies (Thermo Fisher Scientific, Waltham, MA, USA). Near-infrared fluorophore-conjugated secondary antibodies (LI-COR Biosciences, Lincoln, NE, U.S.A). Immunoblotting was performed as described previously [5]. Following secondary antibody incubation, membranes were washed thrice with 1X TBST and imaged using either ChemiDoc (Cambridge Bioscience, Cambridge, United Kingdom) or a Li-Cor Odyssey CLx imaging system (LI-COR Biosciences).

For the immunocytochemical assay, cells were processed as previously described [5]. Cells were then mounted using Vectashield mounting medium (Vector Laboratories Inc., Burlingame, CA, USA) with DAPI and visualized using Leica SP-8 confocal laser scanning microscope (Leica microsystems, Wetzlar, Germany) at ×63 magnification.

### Plasmids

Full-length and C-terminal deletion constructs of ARID1B were used as described previously [5]. Wild-type and catalytically inactive mutants of WWP1 (C890S) and WWP2 (C838A), wild-type and lysine-mutated Ubiquitin constructs, and shRNA constructs for WWP2 were kind gifts from Dr. M. S. Reddy (BRIC**-**CDFD, Hyderabad, India). Full length (Wild-type (WT)), as well as several domain deletions (C2 deletion (ΔC2), C2 plus all four WW domain deletions (ΔC2+ΔWW), and HECT domain deletion (ΔHECT) of WWP2 were generated. ARID1B PPxY deletion mutants were generated by site-directed mutagenesis using specific primers (PPxY #1 forward: 5’ CCTCAGCAGCAGATGGGACAGCAAGGTGTG 3’, reverse: 5’ CACACCTTGCTGTCCCATCTGCTGCTGAGG 3’; PPxY #2 forward: 5’ GGTAACTACTCCAGAAGTGGGGTGCCCAGT 3’, reverse: 5’ ACTGGGCACCCCACTTCTGGAGTAGTTACC 3’). All constructs were confirmed by sequencing.

### Affinity purification and Mass spectrometry

For SFB (Streptavidin-binding protein, Flag, and S-peptide) tagged proteins, Streptavidin-sepharose bead (Cat# 17-511301; GE Healthcare Bio-Science, Uppsala, Sweden) and for Halo tagged proteins, HaloTag bead (Halo pulldown and labeling kit, Cat# G6500; Promega Corporations) based affinity pulldown and antibody-based immunoprecipitations (Protein A Mag Sepharose, Cat # 28944006, Cytiva, United States) were performed as described previously [5,19].

Liquid Chromatography Mass Spectrometry (LC-MS) (done twice - LMO1 and LMO2 (in duplicate)) was performed as per standard protocol. Briefly, HaloTag pull-down assay was carried out using the HaloTag Mammalian Pull-Down System (Promega Corporations) according to the manufacturer’s protocol. 30 hours post transfection by Halo-vector or Halo-ARID1B, HEK293T cells were collected, washed, and lysed using Halo lysis buffer. Equilibrated HaloLink Resin was mixed with the lysate, incubated, washed, and eluted with SDS elution buffer. The eluted samples were analyzed by SDS-PAGE and detected by staining with Coomassie Brilliant Blue R250. The protein bands were excised from the gels and subjected to LC-MS (Taplin Biological Mass Spectrometry Facility, Harvard Medical School, Boston, MA 02115, USA). Interactors present in ARID1B interactome with >1.5-fold change in unique peptide numbers compared to vector control were selected for further analysis.

Mass spectrometric analysis of the ubiquitinated ARID1B protein was performed as per standard protocol. Briefly, 24 hours post co-transfection by hemagglutinin (HA)-Ubiquitin and SFB-ARID1B in presence or absence of Myc-WWP2-WT in biological triplicates, HEK293T cells were lysed under denaturing conditions (1% SDS and heat activated denaturation followed by NETN lysis buffer). Streptavidin bead-based pull-down was performed. The analytes were separated by SDS-PAGE and stained with Coomassie dye. Excised protein bands were subjected to mass spectrometry at the proteomics facility of BRIC-National Centre for Cell Science (BRIC-NCCS), Pune, Maharashtra, India, where proteins were eluted from the gel, reduced with DTT, alkylated with IAA, and digested by trypsin at a ratio of 1:50. The peptide samples were concentrated, desalted using C18 ZipTips (Millipore, Burlington, Massachusetts, United States), and reconstituted in LC-MS grade water with 0.1% formic acid. The peptide concentrations were estimated using NanoDrop One (Thermo Scientific, Waltham, MA, USA). 1μg peptides mix of sample was analyzed by Orbitrap Fusion mass spectrometer coupled to EASY- Spray nano flow column (50 cm×75 μm ID, PepMap C18; Thermo Scientific). Protein identification was performed by Proteome Discoverer (version 2.2, Thermo Scientific) through the Sequest HT and MS Amanda databases with FDR < 0.01. Enriched ubiquitinated tryptic peptides were selected based on the abundance ratio of > 1.5 in ‘+WWP2’ samples compared to ‘-WWP2’ samples.

### Cycloheximide chase assay

HEK293T cells were transfected with various combinations of plasmids and cycloheximide (Cat # 01810, Sigma-Aldrich, St. Louis, MO, USA) (50 μg/ml) was added 24 h post-transfection. Cells were harvested at the indicated time points and protein levels were determined by immunoblotting. Densitometry analysis of bands was done using Fiji software.

### Ubiquitination assay

HEK293T or HCT116 cells were transfected with different combinations of wild-type and mutant ubiquitin, each containing one wild-type lysine, along with SFB tagged ARID1B. At 24 h post-transfection, cells were treated with MG132 (Cat # C2211, Sigma-Aldrich) (10 μM) for 6 h. After harvesting, cells were denatured by 1% SDS followed by 10 min of boiling at 95°C. The lysate was diluted with NETN lysis buffer (20 mM Tris-HCl pH 8.0, 100 mM NaCl, 1 mM EDTA, and 0.5% Nonidet P-40) to bring the SDS concentration to 0.1%. The solution was sonicated using the QSONICA Q500 (Qsonica L.L.C, Newtown, CT, USA) at 10% amplitude for 4 x 5s pulses and centrifuged at 14000 g for 10 min. The supernatant was added to pre-equilibrated streptavidin sepharose beads and incubated for 2 hours at 4°C. The analysis of ubiquitination was carried out by immunoblotting with anti-HA tag antibody.

### Mice tumor xenografts

All xenograft experiments were conducted with prior approval from the Institutional Animal Ethics Committee of the BRIC-Centre for DNA Fingerprinting and Diagnostics (BRIC-CDFD; proposal no. EAF/MDB/12/2025). Female Foxn1^−/−^ nude mice (7-8 weeks old; n = 6) were randomly assigned to experimental groups and subcutaneously inoculated with HCT116 cells exhibiting either ARID1B or ARID1B+WWP2 shRNA mediated knockdown with non-targeting shRNA as the control. Tumor dimensions were measured every 7 days using a vernier caliper, and volumes were calculated using the formula V = (W² × L)/2, where W and L denote tumor width and length, respectively. Five weeks after inoculation, mice were euthanized by Carbon dioxide inhalation, and tumors were excised for determination of weight and volume.

### Statistical analysis

All data obtained from three independent experiments were represented as mean ±s.d. Student’s t-test (unpaired) or two-way ANOVA test was used to determine the statistical significance of all experiments.

### Author Contributions

**PH:** Conceptualization, Methodology, Investigation, Formal analysis, Validation, Data Curation, Visualization, Writing- Original draft, Writing - Review & Editing. **SB:** Methodology, Investigation, Formal analysis, Validation, Visualization, Writing - Review & Editing. **MDB:** Conceptualization, Resources, Formal analysis, Supervision, Project administration, Funding acquisition, Writing- Original draft, Writing - Review & Editing.

## Acknowledgements

We thank Dr. M. S. Reddy, BRIC-Centre for DNA Fingerprinting and Diagnostics (BRIC-CDFD), Hyderabad, India, for providing wild-type and catalytically inactive mutants of WWP1 and WWP2, wild-type and lysine-mutated Ubiquitin constructs, and shRNA constructs for WWP2. HEK293T cells were kind gift from Dr. Rashna Bhandari, BRIC-Centre for DNA Fingerprinting and Diagnostics (BRIC-CDFD), Hyderabad, India. We thank Dr. Srinivas Animireddy (Fellow-E, BRIC-inStem, Bangalore, Karnataka, India) for initial ARID1B interactome data generation (LMO1) and for identifying possible interaction of ARID1B with WWP2. We acknowledge the Sophisticated Equipment Facility at CDFD for assistance with confocal microscopy and Sanger sequencing, and the Experimental Animal Facility for support with the nude mice xenograft experiments. Graphical illustrations were generated using BioRender (https://www.biorender.com/).

## Data Availability Statement

The mass spectrometry data from this study are available on MassIVE (https://massive.ucsd.edu), a Proteome Xchange Consortium member, and can be accessed through the dataset identifier MSV000101989.

## Abbreviations

SWI/SNF: SWItch/Sucrose Non-Fermentable
BAF BRG1/: Brm Associated Factor
cBAF: Canonical BAF
PBAF: Polybromo BAF
ncBAF: Non-canonical BAF
GBAF: GLTSCR1-like containing BAF
ARM: Armadillo
NLS: Nuclear Localisation Signal
IDR: Intrinsically Disordered Region
RING: Really Interesting Gene
HECT: Homologous to the E6-associated protein carboxyl domain
RBR: RING-between-RING
WWP1/2: WW domain-containing E3 ubiquitin protein ligase 1/2
GO: Gene Ontology
ARID1B: AT-rich interaction domain protein 1B
Ub: Ubiquitin
HA: Hemagglutinin
SFB: Streptavidin-binding protein, Flag, and S-peptide
GFP: Green Fluorescent protein
LC-MS/MS: Liquid Chromatography-Tandem Mass Spectrometry
Dvl2: Dishevelled segment polarity protein 2
PTEN: Phosphatase and TENsin homolog deleted on chromosome 10
SHP-1: Src homology region 2 domain-containing phosphatase-1
Cul2: Cullin-2
EloC: Elongin C
Roc1: Regulator of Cullins-1
HACE1: HECT domain and ankyrin repeat-containing E3 ubiquitin-protein ligase1

---

**Supp. Fig. 1 (related to Fig. 1):**
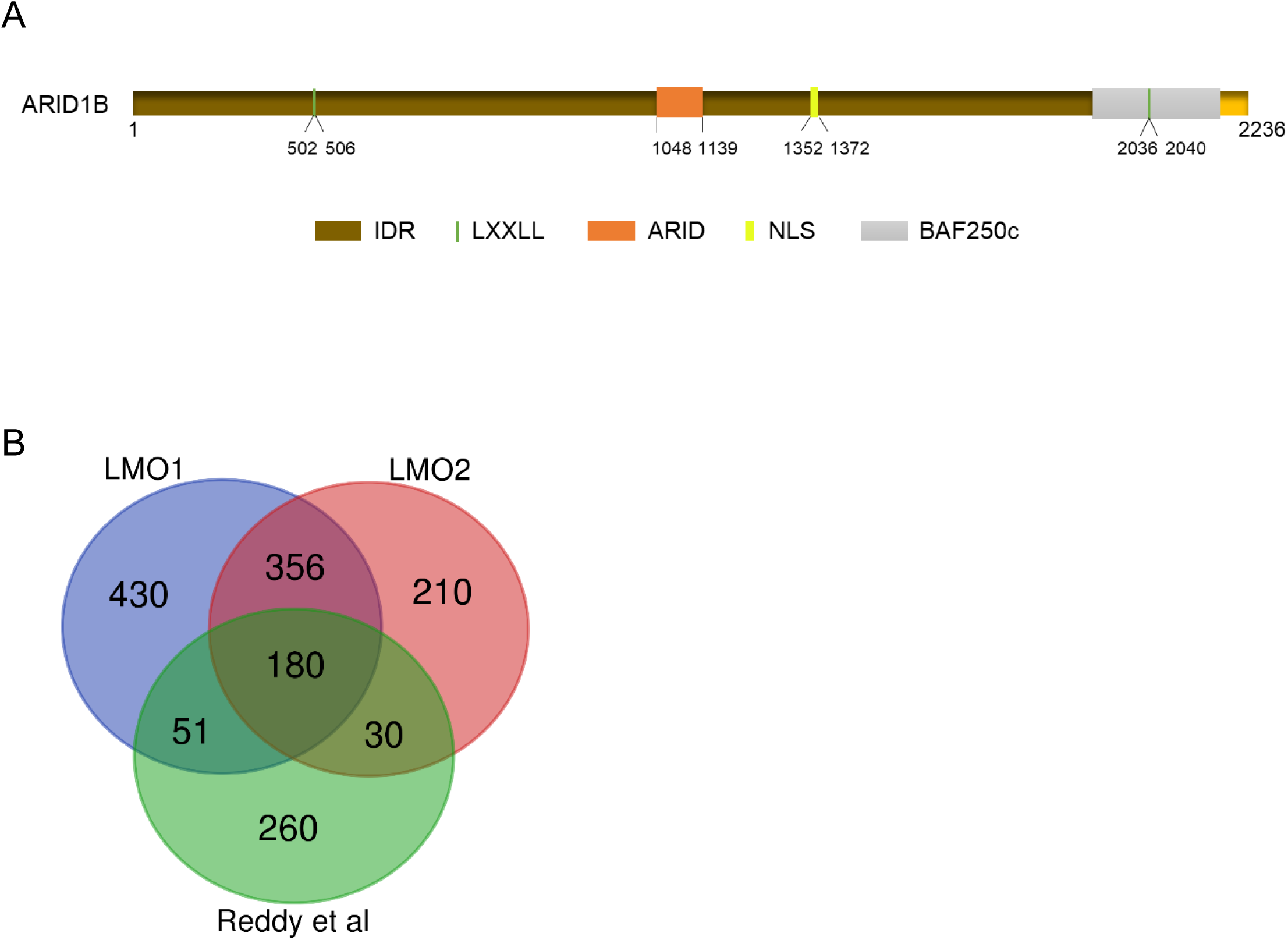
Schematic representation of ARID1B domains (A). The major domains are shown in distinct colors: the intrinsically disordered region (IDR) in brown, the LXXLL motif as a graphical bar, the ARID domain in orange, the nuclear localization signal (NLS) in yellow, and the BAF250_C domain in grey. Common interacting proteins identified in multiple ARID1B interactome screens (B).

**Supp. Fig. 2 (related to Fig. 2):**
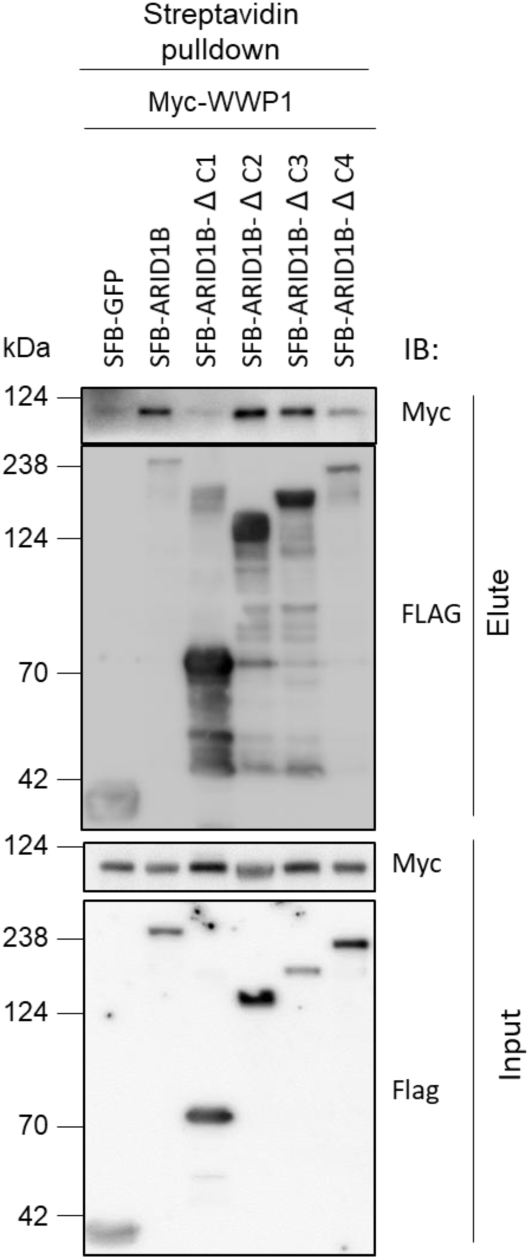
Interaction between Myc-WWP1 and full-length and various C-terminal deletion mutants of SFB-tagged ARID1B (B).

**Supp. Fig. 3 (related to Fig. 3):**
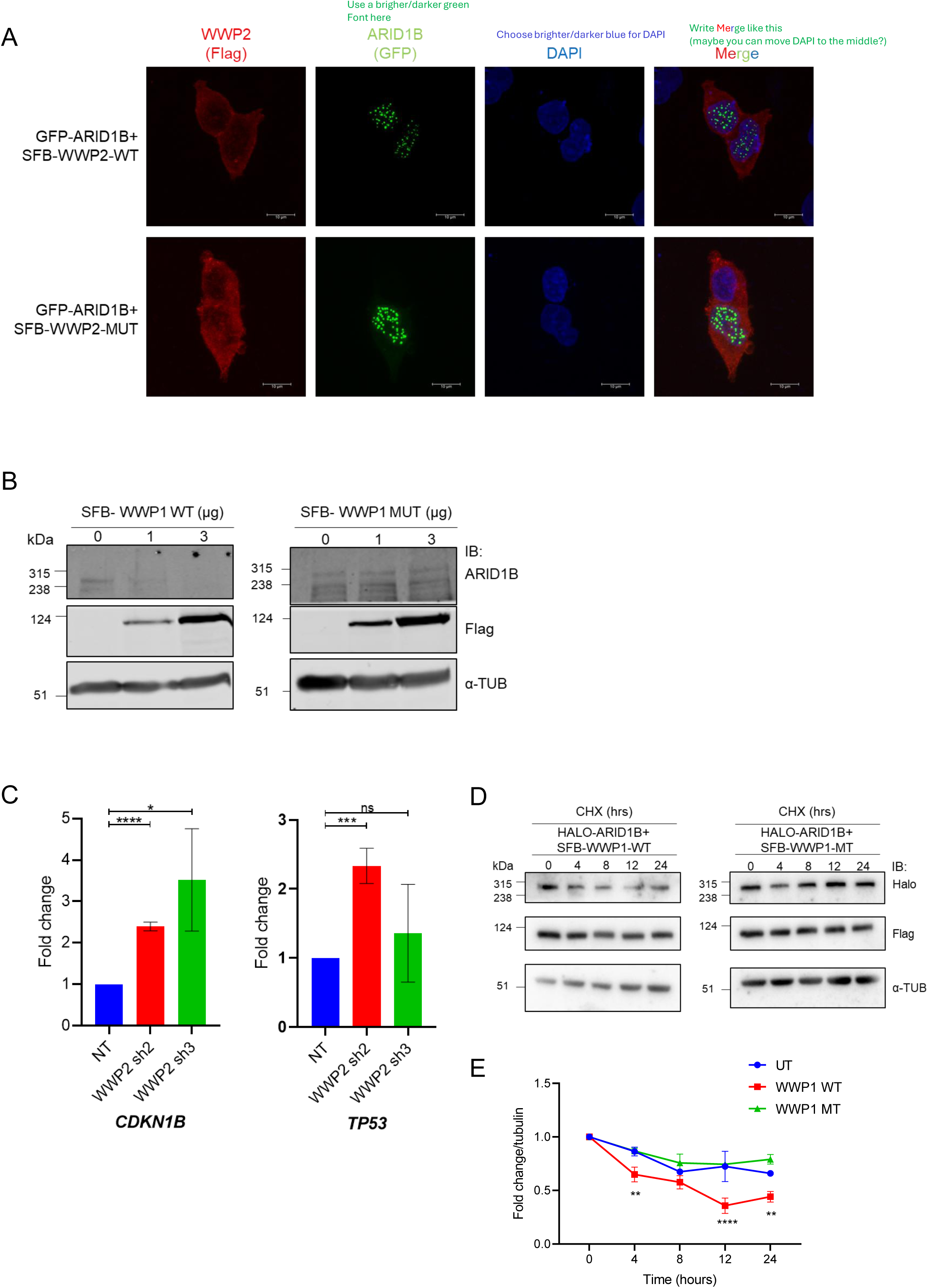
: Intracellular localisation of GFP-ARID1B was assessed upon ectopic expression of either wild-type or catalytically inactive mutant of SFB-WWP2 (A). Assessment of endogenous ARID1B levels when exposed to increasing amounts of ectopically expressed wild-type or mutant WWP1 (B). Evaluation of ARID1B target gene transcript levels upon downregulation of WWP2 in HCT116 cells (C). Evaluation of ARID1B protein stability by cycloheximide chase assay in the presence or absence of wild-type or mutant WWP1 (D); quantification is shown in panel E. WT, wild-type; MT, mutant; UT, untransfected. Quantifications (panel C) are from three independent experiments. All experiments were performed at least in three biological replicates; representative images are shown. All statistical analyses were performed using two-way ANOVA (N=3), \**P*<0.05; \*\**P*<0.01; \*\*\**P*<0.001; \*\*\*\**P*<0.0001

**Supp. Fig. 4 (related to Fig. 4):**
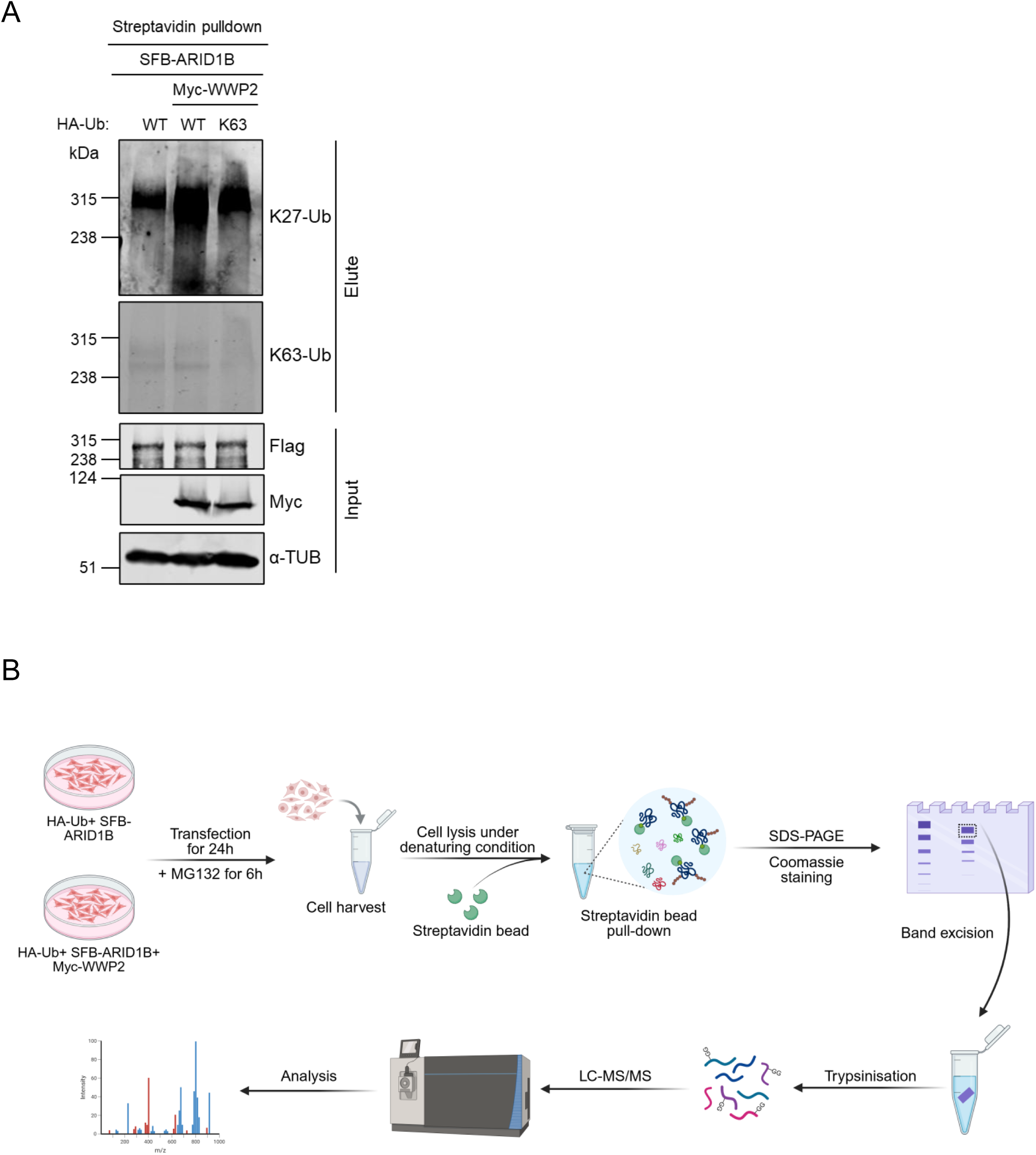
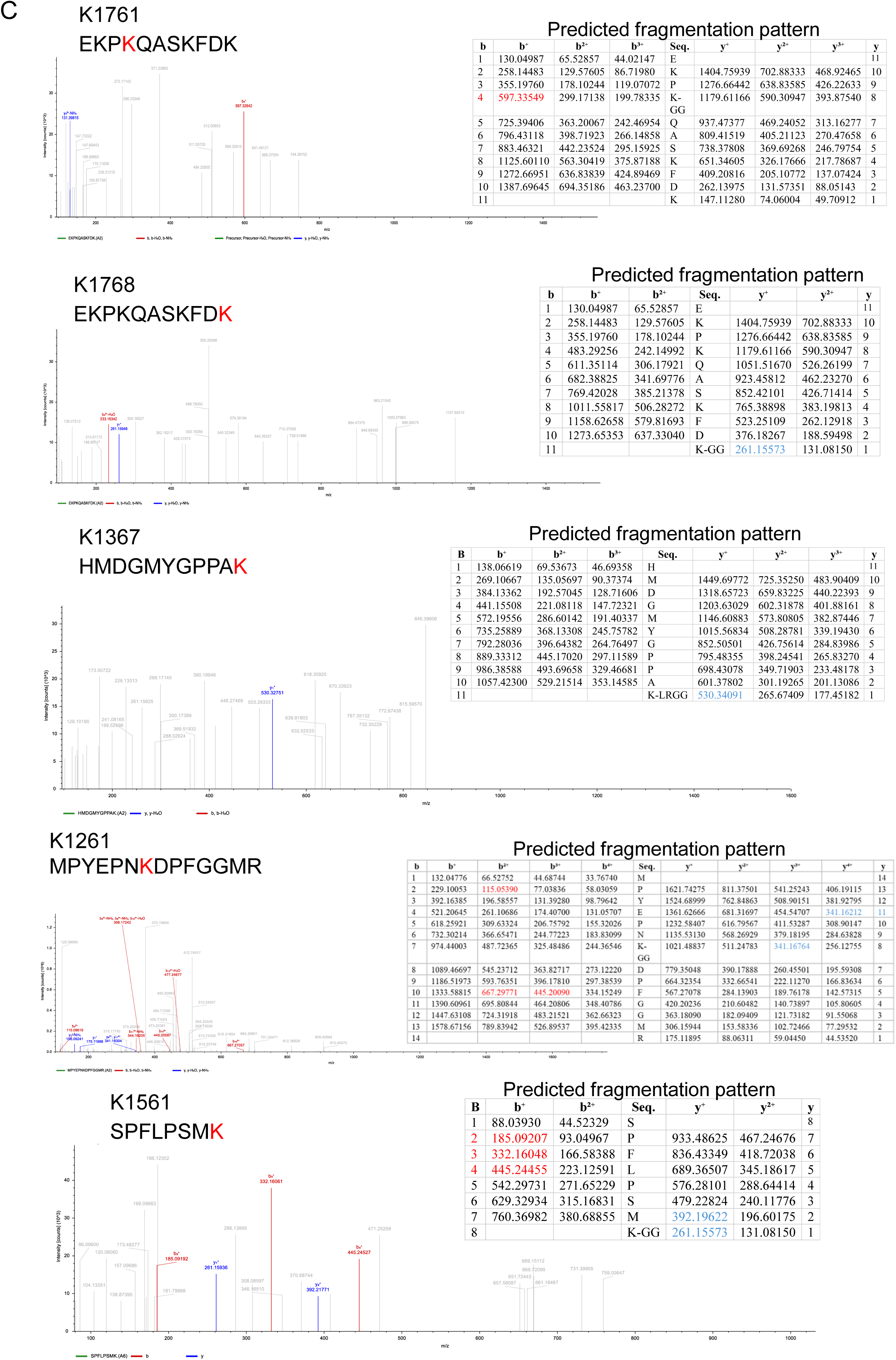
Analysis of linkage-specific polyubiquitin chain elongation of ARID1B in the presence of WWP2, upon immunoblotting with endogenous K27-linkage and K63-linkage specific ubiquitin antibodies (A). Schematic diagram depicting the methodology followed in identifying enriched ubiquitinated peptides of ARID1B in presence of WWP2 using LC-MS/MS based analysis (B). Representative LC-MS/MS spectra of the enriched ubiquitinated peptides of ARID1B upon overexpression of WWP2. The sequence of the tryptic fragment is shown above each spectrum, and the modified residue is highlighted in ‘red’ font. The masses of the b ions (red) and y ions (blue) detected in the peptide fragmentation pattern are shown in the table to the right (C).

**Supp Fig. 5 (related to Fig. 5):**
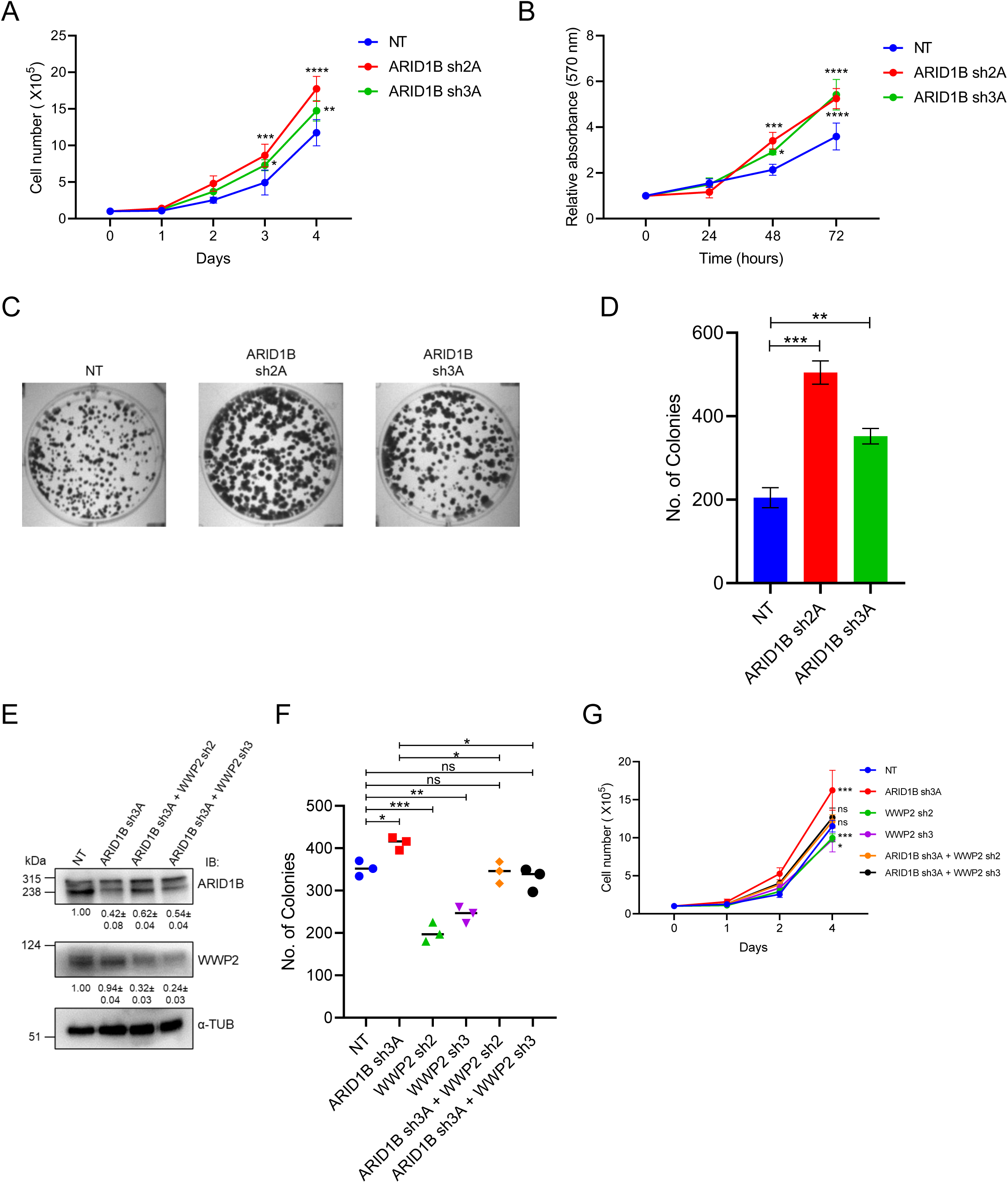
Cell growth (A), viability (B), and colony formation (C, D) assays performed in HCT116 cells following ARID1B knockdown. shRNA mediated knockdown of ARID1B (sh3A) and subsequently of WWP2 (sh2 and sh3) in HCT116 colorectal cancer cells (E). colony formation (F), and cell growth (G) assays were performed with various combinations of downregulation of ARID1B and WWP2, as indicated. All experiments were performed at least in three biological replicates; representative images are shown. All statistical analyses were performed using either unpaired Student’s T test or two-way ANOVA (N=3), \**P*<0.05; \*\**P*<0.01; \*\*\**P*<0.001; \*\*\*\**P*<0.0001

**Supp. Fig. 6:**
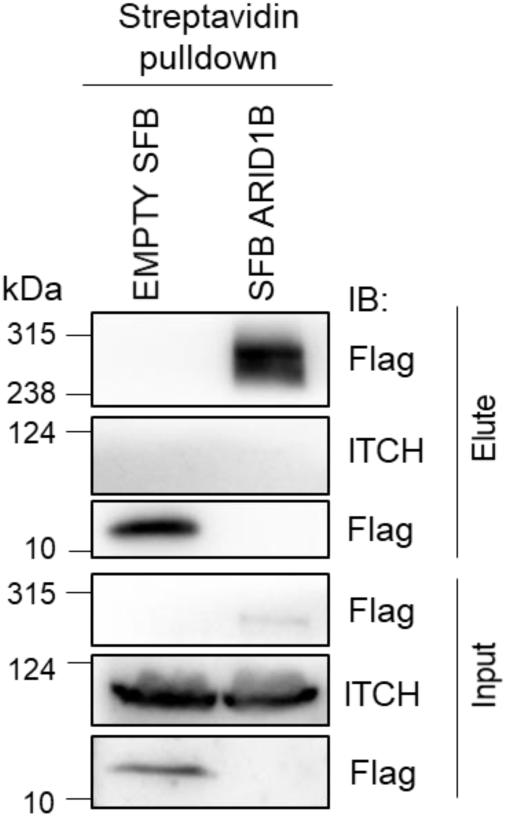
Absence of interaction between ARID1B and ITCH.

**Supp. Table 1:**
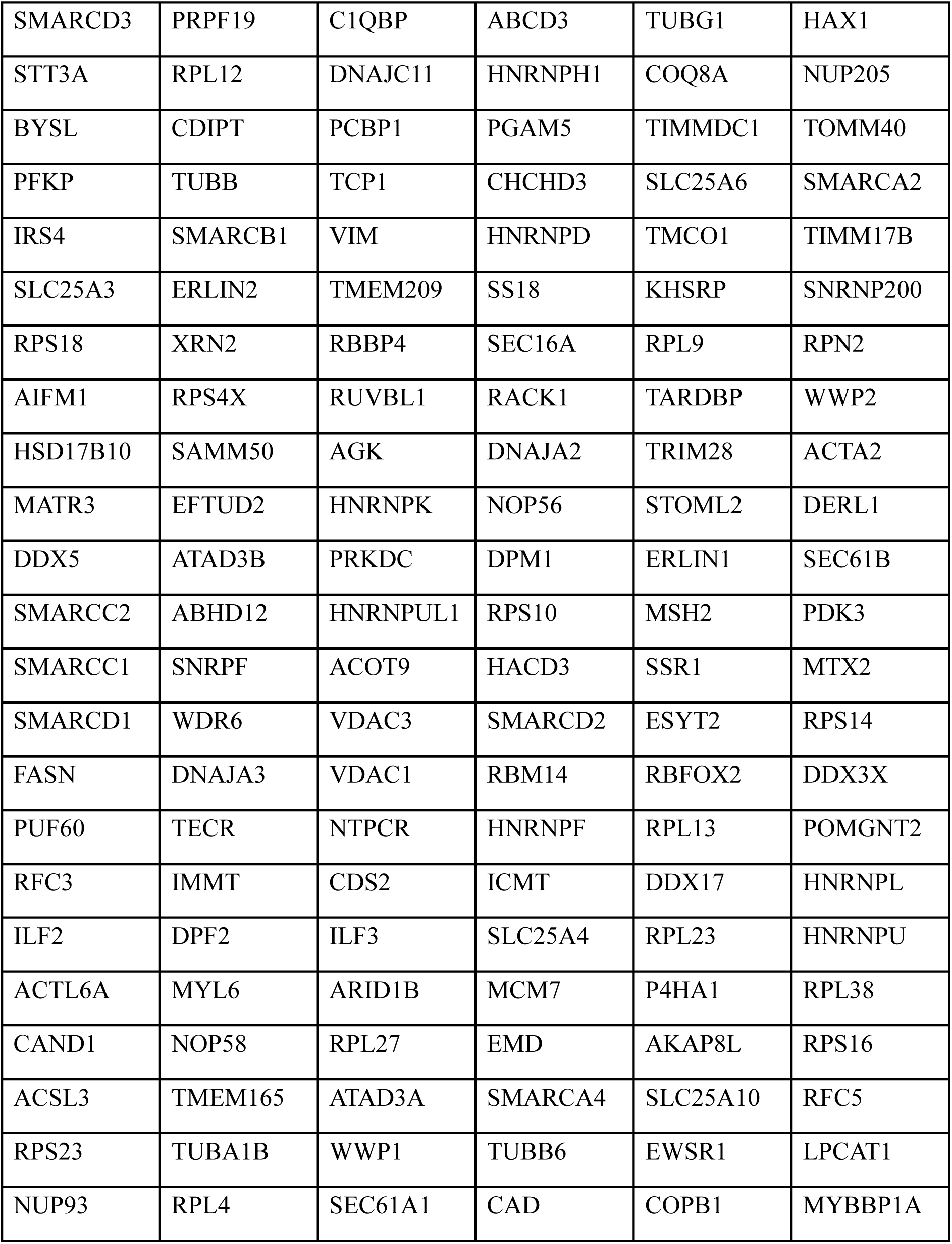
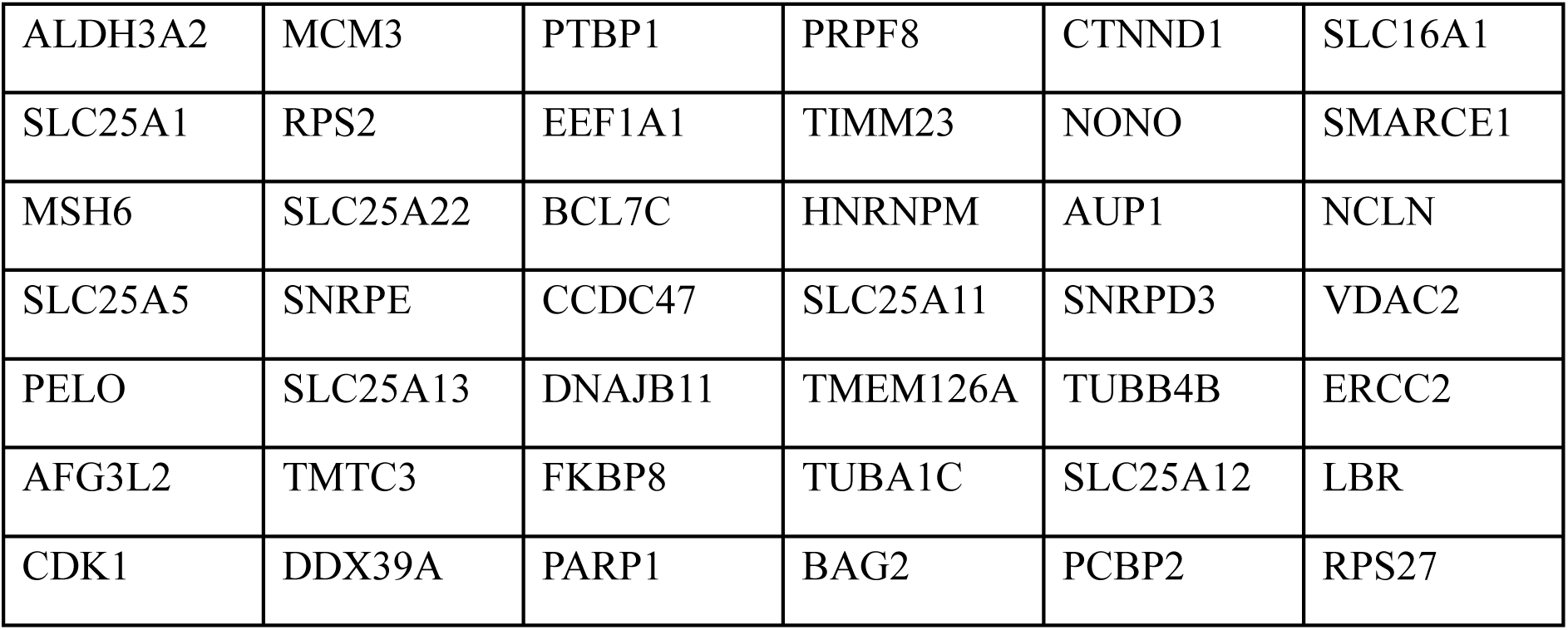
List of common interacting partners of ARID1B obtained from different interactomes.

|  |  |  |  |  |  |
| --- | --- | --- | --- | --- | --- |
| SMARCD3 | PRPF19 | C1QBP | ABCD3 | TUBG1 | HAX1 |
| STT3A | RPL12 | DNAJC11 | HNRNPH1 | COQ8A | NUP205 |
| BYSL | CDIPT | PCBP1 | PGAM5 | TIMMDC1 | TOMM40 |
| PFKP | TUBB | TCP1 | CHCHD3 | SLC25A6 | SMARCA2 |
| IRS4 | SMARCB1 | VIM | HNRNPD | TMCO1 | TIMM17B |
| SLC25A3 | ERLIN2 | TMEM209 | SS18 | KHSRP | SNRNP200 |
| RPS18 | XRN2 | RBBP4 | SEC16A | RPL9 | RPN2 |
| AIFM1 | RPS4X | RUVBL1 | RACK1 | TARDBP | WWP2 |
| HSD17B10 | SAMM50 | AGK | DNAJA2 | TRIM28 | ACTA2 |
| MATR3 | EFTUD2 | HNRNPK | NOP56 | STOML2 | DERL1 |
| DDX5 | ATAD3B | PRKDC | DPM1 | ERLIN1 | SEC61B |
| SMARCC2 | ABHD12 | HNRNPUL1 | RPS10 | MSH2 | PDK3 |
| SMARCC1 | SNRPF | ACOT9 | HACD3 | SSR1 | MTX2 |
| SMARCD1 | WDR6 | VDAC3 | SMARCD2 | ESYT2 | RPS14 |
| FASN | DNAJA3 | VDAC1 | RBM14 | RBFOX2 | DDX3X |
| PUF60 | TECR | NTPCR | HNRNPF | RPL13 | POMGNT2 |
| RFC3 | IMMT | CDS2 | ICMT | DDX17 | HNRNPL |
| ILF2 | DPF2 | ILF3 | SLC25A4 | RPL23 | HNRNPU |
| ACTL6A | MYL6 | ARID1B | MCM7 | P4HA1 | RPL38 |
| CAND1 | NOP58 | RPL27 | EMD | AKAP8L | RPS16 |
| ACSL3 | TMEM165 | ATAD3A | SMARCA4 | SLC25A10 | RFC5 |
| RPS23 | TUBA1B | WWP1 | TUBB6 | EWSR1 | LPCAT1 |
| NUP93 | RPL4 | SEC61A1 | CAD | COPB1 | MYBBP1A |
| ALDH3A2 | MCM3 | PTBP1 | PRPF8 | CTNND1 | SLC16A1 |
| SLC25A1 | RPS2 | EEF1A1 | TIMM23 | NONO | SMARCE1 |
| MSH6 | SLC25A22 | BCL7C | HNRNPM | AUP1 | NCLN |
| SLC25A5 | SNRPE | CCDC47 | SLC25A11 | SNRPD3 | VDAC2 |
| PELO | SLC25A13 | DNAJB11 | TMEM126A | TUBB4B | ERCC2 |
| AFG3L2 | TMTC3 | FKBP8 | TUBA1C | SLC25A12 | LBR |
| CDK1 | DDX39A | PARP1 | BAG2 | PCBP2 | RPS27 |

